# Fibroblast-specific inflammasome activation predisposes to atrial fibrillation

**DOI:** 10.1101/2023.05.18.541326

**Authors:** Luge Li, Cristian Coarfa, Yue Yuan, Issam Abu-Taha, Xiaolei Wang, Jia Song, Amrit Koirala, Sandra L Grimm, Markus Kamler, Lisa K Mullany, Michelle Tallquist, Stanley Nattel, Dobromir Dobrev, Na Li

## Abstract

**Background:** Recent work has shown that the NLR-family-pyrin-domain-containing 3 (NLRP3) inflammasome is expressed in cardiomyocytes and when specifically activated causes atrial electrical remodeling and arrhythmogenicity. Whether the NLRP3-inflammasome system is functionally important in cardiac fibroblasts (FBs) remains controversial. In this study, we sought to uncover the potential contribution of FB NLRP3-inflammasome signaling to the control of cardiac function and arrhythmogenesis.

**Methods:** Digital-PCR was performed to determine the expression of NLRP3-pathway components in FBs isolated from human biopsy samples of AF and sinus rhythm patients. NLRP3-system protein expression was determined by immunoblotting in atria of canines with electrically maintained AF. Using the inducible, resident fibroblast (FB)-specific Tcf21-promoter-Cre system (Tcf21iCre as control), we established a FB-specific knockin (FB-KI) mouse model with FB-restricted expression of constitutively active NLRP3. Cardiac function and arrhythmia susceptibility in mice were assessed by echocardiography, programmed electrical stimulation, and optical mapping studies.

**Results:** NLRP3 and IL1B were upregulated in atrial FBs of patients with persistent AF. Protein levels of NLRP3, ASC, and pro-Interleukin-1β were increased in atrial FBs of a canine AF model. Compared with the control mice, FB-KI mice exhibited enlarged left atria (LA) and reduced LA contractility, a common determinant of AF. The FBs from FB-KI mice were more transdifferentiated, migratory, and proliferative compared to the FBs from control mice. FB-KI mice showed increased cardiac fibrosis, atrial gap junction remodeling, and reduced conduction velocity, along with increased AF susceptibility. These phenotypic changes were supported by single nuclei (sn)RNA-seq analysis, which revealed enhanced extracellular matrix remodeling, impaired communication among cardiomyocytes, and altered metabolic pathways across multiple cell types.

**Conclusions:** Our results show that the FB-restricted activation of the NLRP3-inflammasome system leads to fibrosis, atrial cardiomyopathy, and AF. Activation of NLRP3-inflammasome in resident FBs exhibits cell-autonomous function by increasing the activity of cardiac FBs, fibrosis, and connexin remodeling. This study establishes the NLRP3-inflammasome as a novel FB-signaling pathway contributing to AF pathogenesis.

## INTRODUCTION

Atrial fibrillation (AF), the most cardiac common arrhythmia, is associated with increases in morbidity and mortality^1^. AF typically requires ectopic (triggered) firing for induction and a reentrant substrate for maintenance^2^. The reentrant substrate may result from reduced atrial refractoriness, conduction disturbances, and/or structural changes including fibrosis and dilatation^3^. It is increasingly recognized that atrial cardiomyopathy, which manifests as fibrosis, reduced left atrial (LA) contractility, and diastolic dysfunction^4-6^ is a major underlying cause of AF. The NLR family pyrin domain containing 3 (NLRP3) inflammasome known to be centrally involved in inflammatory signaling, is responsible for the maturation of caspase-1 (Casp1) and interleukin-1β (IL-1β)^7, 8^. Previous work has shown that cardiomyocyte (CM)-specific NLRP3 inflammasome activation promotes atrial ectopic activity and electrical remodeling that cause AF development ^9-11^. It is also known that NLRP3 inflammasomes exist in cardiac fibroblasts (FBs) ^12, 13^; however, the contribution of FB NLRP3 inflammasome signaling to atrial remodeling and arrhythmogenesis is poorly understood. Because FBs play an important role in pathologies of AF and atrial cardiomyopathy, we hypothesized that FB-specific activation of the NLRP3 inflammasome might promote the development of an AF-promoting cardiomyopathy. Using a FB-specific knockin (FB-KI) mouse model that expresses constitutively active NLRP3 in resident FBs, we show here that the activation of FB NLRP3 inflammasomes promotes the development of atrial cardiomyopathy, abnormal conduction, cardiac fibrosis, and extracellular matrix remodeling, thereby increasing the predisposition to AF.

## METHODS

### Human atrial samples

Right atrial appendage (RAA) samples were collected from patients undergoing open-heart surgery for coronary bypass grafting and/or valve replacement. Six control patients with NSR and six with persistent AF (perAF, >6-month duration) provided informed consent prior to surgery. All experimental protocols were approved by the Human Ethics Committee of the Medical Faculty of the University Duisburg-Essen (approval number AZ:12-5268-BO) and were performed in accordance with the Declaration of Helsinki.

### Canine model of AF

Studies involving dogs were approved by the Animal Research Ethics Committee of the Montreal Heart Institute (protocol 2018-47-12) and followed Canadian Council on Animal Care Guidelines. Upon arbitrary assignment, 13 mongrel adult dogs were implanted with a right-atrial pacemaker. AF was maintained for 1 week by right-atrial pacing at 600 bpm. Dogs in the control (Ctl) group received a turned-off pacemaker ^14-16^. Left-atrial FBs were isolated as previously described ^17^.

### FB-specific NLRP3 knockin mouse model

All studies involving mice were performed according to protocols approved by the Institutional Animal Care and Use Committee at Baylor College of Medicine and conformed to the *Guide for the Care and Use of Laboratory Animals* published by the U.S. National Institutes of Health. To yield mice with FB-specific expression of constitutively active NLRP3, *Tcf21*^*iCre*^ mice were crossbred to *Nlrp3*^*neoR-A350V*^ mice (*Tcf21*^*iCre*^*:Nlrp3*^*A350V/A350V*^, FB-KI) ^18, 19^. To activate Cre recombinase, FB-KI mice were injected with tamoxifen (50mg/kg, i.p.) for 5 days. *Tcf21*^*Cre*^*:Nlrp3*^*WT/WT*^ mice injected with tamoxifen were used as Ctl. The offspring of this cross maintained expected Mendelian ratios. To avoid the potential interference with Cre activity due to the estrogen receptor, only male mice were used in this study.

### Digital PCR in human atrial FBs

Total RNA was extracted from the isolated atrial FBs using the RNeasy Mini Kit (Qiagen, Hilden, Germany) and 500 ng RNA was transcribed into cDNA using a reverse transcription kit (Applied Biosystems; Thermo Fisher Scientific, Waltham, MA). Digital PCR (dPCR) was carried out with TaqMan probes from Thermo Fisher Scientific.

### *In vivo* electrophysiology

To assess AF-inducibility, programmed electrical stimulation was performed 1-month post tamoxifen injection in Ctl and FB-KI mice as described previously ^9^. Each mouse was subjected to 3 sequences of rapid atrial pacing to induce AF. When an animal developed atrial tachycardia/AF episodes longer than 1-second at least 2 times, it was considered AF-positive. AF-inducibility was calculated as the percentage of AF-positive mice over the total number of mice studied.

### Isolation and culture of primary mouse cardiac FBs

Adult FBs were isolated from whole hearts of Ctl and FB-KI mice. Briefly, heart was dissected from the anesthetized mouse, placed in cold KH buffer to remove excess blood, and cut into small pieces. Afterwards, 5 mL of digestion cocktail containing 8.75 μL Dase I, 90 μL HEPES, 250 μL collagenase and 10 mL HBSS, was added to the sample. After incubation at 37°C for 45 min with gentle rocking, the supernatant was resuspended with KH buffer, followed by filtration with cell strainer (gap size: 40 μm) and centrifugation (400x g, 4°C, 10 min). The pellet was resuspended with 5 mL RBC lysis buffer and incubated at room temperature for 2 min and centrifuged (400x g, 4°C, 10 min) to remove the supernatant. The pellet was suspended with culture media and plated in culture dish. 2 -4 hours after plating, the supernatant was removed. The cells were washed with media and cultured for 24 -48 hours prior to the following functional assays.

### Migration assay

FBs at 3×10^5^ were placed in 12-well plates and incubated with medium overnight. A scratch was made with the 200 μL pipette tip down the middle well and cells were washed with medium. The cells were imaged and culture medium without FBS was changed every 2 hours for 24 hours.

### Gel contraction assay

Gels were prepared containing 75,000 FBs, 150 μL collagen I, 150 μL DMEM with 20 mM HEPES, and 44 mM NaHCO_3_, and then 310 μL compound was plated in 24-well plate and incubated at 37°C for 25 min. A 28G needle was used to dislodge gel and 600 μL medium with and without TGF-β1 (1 ng/mL) was added. The FB/gel mixtures were then incubated at 37 °C for 24 hours, followed by imaging.

### Edu staining

FBs were placed at 50% confluency in 6-well plates and incubated with EdU (10 μM) for 4 hours, followed by fixation with 4% PFA for 15 min at room temperature. Hoechst 33342 (3 μg/mL) was used to stain nuclei. Fluorescent images were acquired with a fluorescent microscope. The number of EdU+ cells and the total number of cells in five randomly selected microscopic visual fields were counted with Image J.

### Single nuclei RNA-Seq

Single nuclei (sn) were isolated according to previous methods^20^. The snRNA-Seq libraries were generated using the 10x Genomics Chromium Single Cell 3’ v3 reagent kit and the sequencing was performed by Novogene. Sequencing reads (GEO accession: GSE218250) were mapped to the mouse reference genome using the CellRanger software from 10x Genomics. Basic normalization and filtering of low-quality cells or low expressed genes was performed using Scanpy^21^. Doublets were detected using the Scrublet software^22^. Gene expression was converted to Uniform Manifold Approximation and Projection (UMAP) and clusters were detected using Scanpy^21^. Cell types were annotated using cell markers of murine heart previously reported^20, 23^. Differentially expressed genes were determined using the Seurat R package^24^ FindMarkers function, with significance achieved for FDR<0.05 and fold change exceeding 1.5-fold. Enriched pathways at cell-type level were determined using the Gene Set Enrichment Analysis (GSEA) software^25^. Ligand-receptor interaction maps were determined using the CellChat method^26^.

### Statistical analysis

Data were presented as mean ± SEM. Two-tailed Student’s *t*-tests were used to compare data between two groups maintaining normal distributions. Welch correction was applied when variances were significantly different. Mann-Whitney tests were used for datasets with non-normal distribution. Fisher’s exact test was used to compare categorical data. A p-value of less than 0.05 was considered statistically significant.

## RESULTS

### NLRP3 inflammasome signaling is enhanced in cardiac FBs of AF patients and an animal model of AF

To assess the expression of the NLRP3 inflammasome pathway in atrial FBs of AF patients, we utilized the highly sensitive digital PCR platform to determine the mRNA level of *NLRP3, ASC* (encoding apoptosis-associated speck-like protein containing a CARD, ASC) and *IL1B* (encoding IL-1β) in human atrial FBs isolated from patients with NSR or perAF. Digital PCR revealed that levels of *NLRP3* and *IL1B* mRNA were higher in atrial FBs of perAF compared to NSR patients (p=0.018, p=0.0013, **Figure 1A-D**). Protein-expression assay in human samples was impossible because of the limited protein-amount that can be obtained from freshly isolated human atrial FB samples available from operative biopsies and the concerns of phenotypic drift with culture. We assessed the NLRP3-inflammasome protein-expression in an established canine model of electrically maintained AF ^14-16^. Western blotting analysis showed that NLRP3, ASC, and pro-IL-1β protein levels were significantly upregulated in atrial FBs of AF dogs (**Figure 1E-I**). The protein levels of IL-1β were not significantly different between groups (**Figure 1H,J**). These results unveil an association between enhanced expression of the FB NLRP3 inflammasome system and the development of AF.

**Figure 1.**
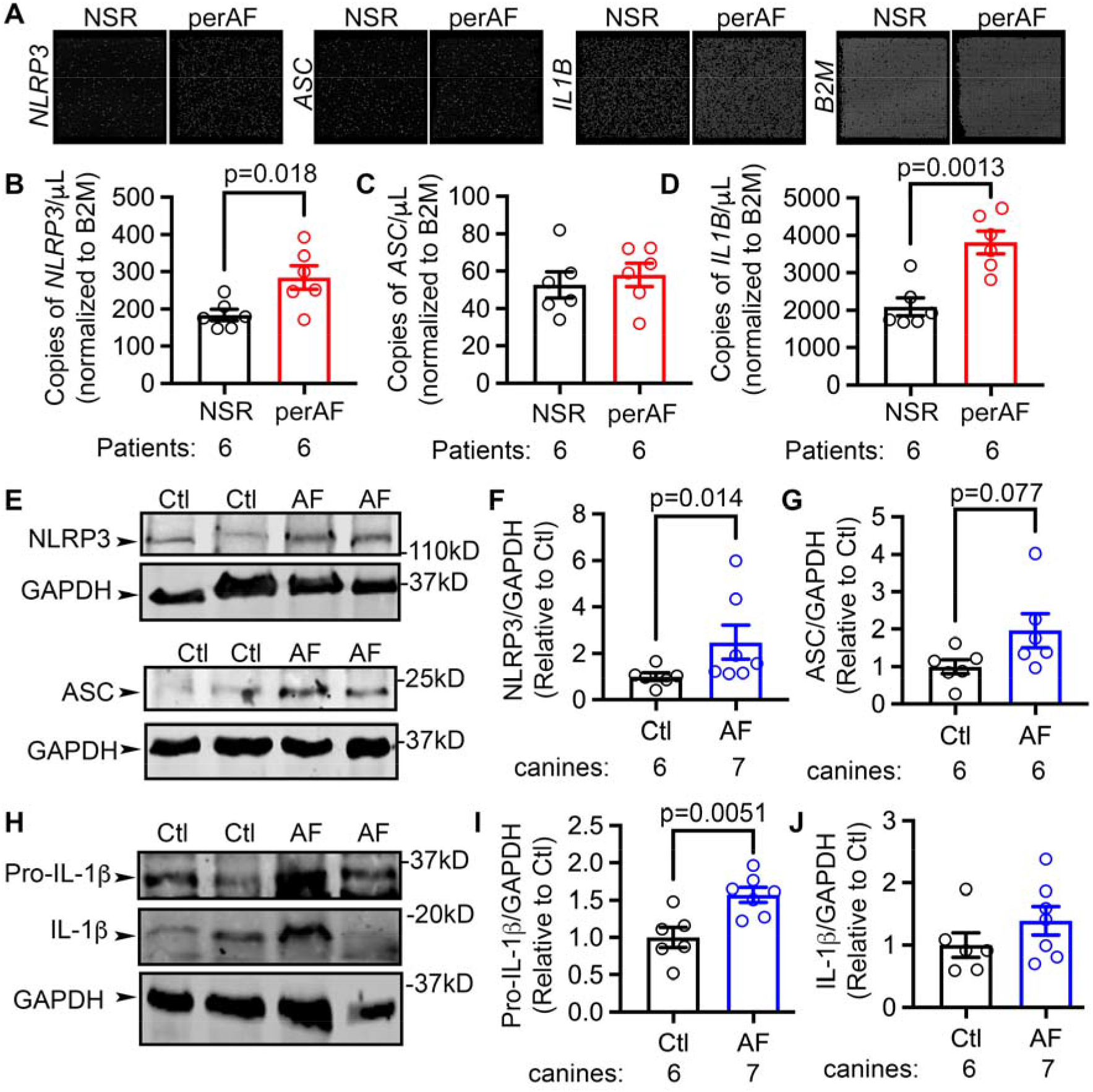
Enhanced NLRP3 inflammasome signaling in FBs of AF patients, canine model of AF, and rat model of RHD. (**A**) Representative digital PCR photos from wells containing 8500 partitions and quantification of *NLRP3* (**B**), *ASC* (**C**), and *IL1B* (**D**) in human atrial FBs isolated from RAA of NSR and perAF patients. (**E, H**) Representative Western blots and quantification of NLRP3 (**F**) and ASC (**G**), and pro- and active IL-1β (**I, J**) in atrial FB-samples of Ctl and AF dogs. p-values were determined with unpaired Student’s t-test (two-tailed) in **B, D, G**, and **I**, and Mann-Whitney test in **F**.

### FB NLRP3-inflammasome activation causes atrial hypocontractility and enhances susceptibility to AF

To elucidate the role of FB NLRP3 inflammasome activation in cardiomyopathy and arrhythmogenesis, we established the FB-specific knockin (FB-KI) mouse model with the FB-restricted expression of constitutively active NLRP3 by crossbreeding *Tcf21*^*iCre*^ mice with *Nlrp3*^*neoR-A350V*^ mice (*Tcf21*^*iCre*^*:Nlrp3*^*A350V/A350V*^, FB-KI) ^18, 19^. 4-weeks post tamoxifen injection (**Figure 2A**), we isolated CMs and cardiac FBs from mouse hearts of Ctl and FB-KI mice. Western blots with the separated FB- and CM-fractions showed that the levels of pro-Casp1 and the mature Casp1 (p20) were increased in the FB-fractions of FB-KI mice and remained unchanged in CM-fractions of FB-KI mice (**Figure 2B-D**), validating the FB-specific activation of NLRP3 inflammasome in FB-KI mice. The enhanced activity of FB NLRP3 inflammasome significantly increased the protein levels of mature IL-1β and IL-18 in atrial tissues of FB-KI mice (**Figure 2E-G**). Of note, the serum level of IL-1β, IL-18 and C-reactive protein (CRP) were comparable between Ctl and FB-KI mice, indicating the absence of systemic inflammation in FB-KI mice. To determine the effect of FB-specific activation of the NLRP3-inflammasome system on atrial function, we first assessed atrial structure by performing echocardiography in Ctl and FB-KI mice at 3 months of age (one month after tamoxifen injection). Long-axis images revealed that the superoinferior (SI) and anteroposterior (AP) dimensions and the area of LA were increased in FB-KI mice (vs Ctl mice, **Figure 3A-D**). M-mode recordings also revealed that LA diameters (LADs, LADd) were increased in FB-KI mice (**Figure 3E-G**). Fractional shortening (FS%), calculated as percentage change of LADs and LADs, was also significantly reduced in FB-KI mice (p=0.0003, **Figure 3H**). These results suggest that FB-KI mice exhibited atrial myopathy, as evidenced by the atrial dilatation and hypocontractility. To determine whether FB-specific activation of NLRP3 can promote atrial arrhythmogenesis, we performed programmed electrical stimulation to induce AF. *In vivo*, we found that FB-KI mice were more susceptible to pacing-induced AF than Ctl littermates (67% vs 20%, p=0.036, **Figure 3I-J**). The duration of the induced AF was also longer in FB-KI mice than in Ctl (p=0.041, **Figure 3K**), pointing to the evolution of an arrhythmogenic substrate. To exclude the potential influence of baroreflex, we also determined the AF inducibility *ex vivo* on the Langendorff-perfused hearts. Consistently, FB-KI mouse hearts subjected to burst pacing (S1-S2, CL:100ms) developed AF more frequently than Ctl mice (85% vs 17%, p=0.030), confirming that activation of FB NLRP3-inflammasome creates a substrate for AF development.

**Figure 2.**
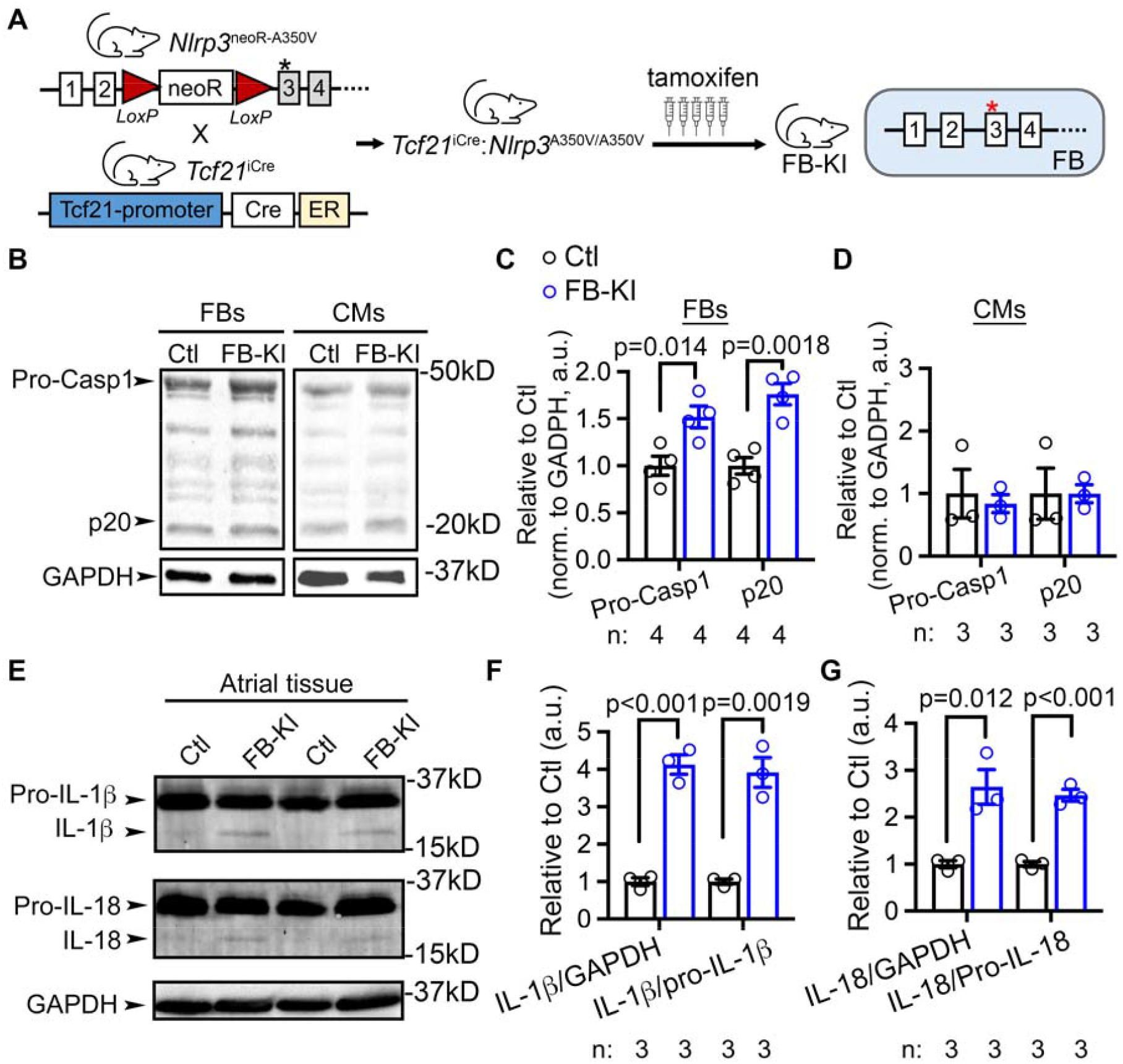
FB-specific activation of NLRP3 in a transgenic murine model. (**A**) Schematic representation of the transgenic approach to establish the inducible *Tcf21*^*iCre*^*;Nlrp3*^*A350V/A350V*^ (FB-KI) mouse model. Tamoxifen injected *Tcf21*^*iCre*^ mice were used as control (Ctl). (**B-D**) Representative Western blots and quantifications of precursor caspase-1 (Pro-Casp1) and cleaved caspase-1 (p20) in separated FB- and CM-fractions of Ctl and FB-KI mice. (**E-G**) Representative Western blots and quantifications of precursor and active IL-1β and IL-18 in atrial tissues of Ctl and FB-KI mice. p-values were determined with unpaired Student’s t-test (two-tailed) in **C, F**, and **G**.

**Figure 3.**
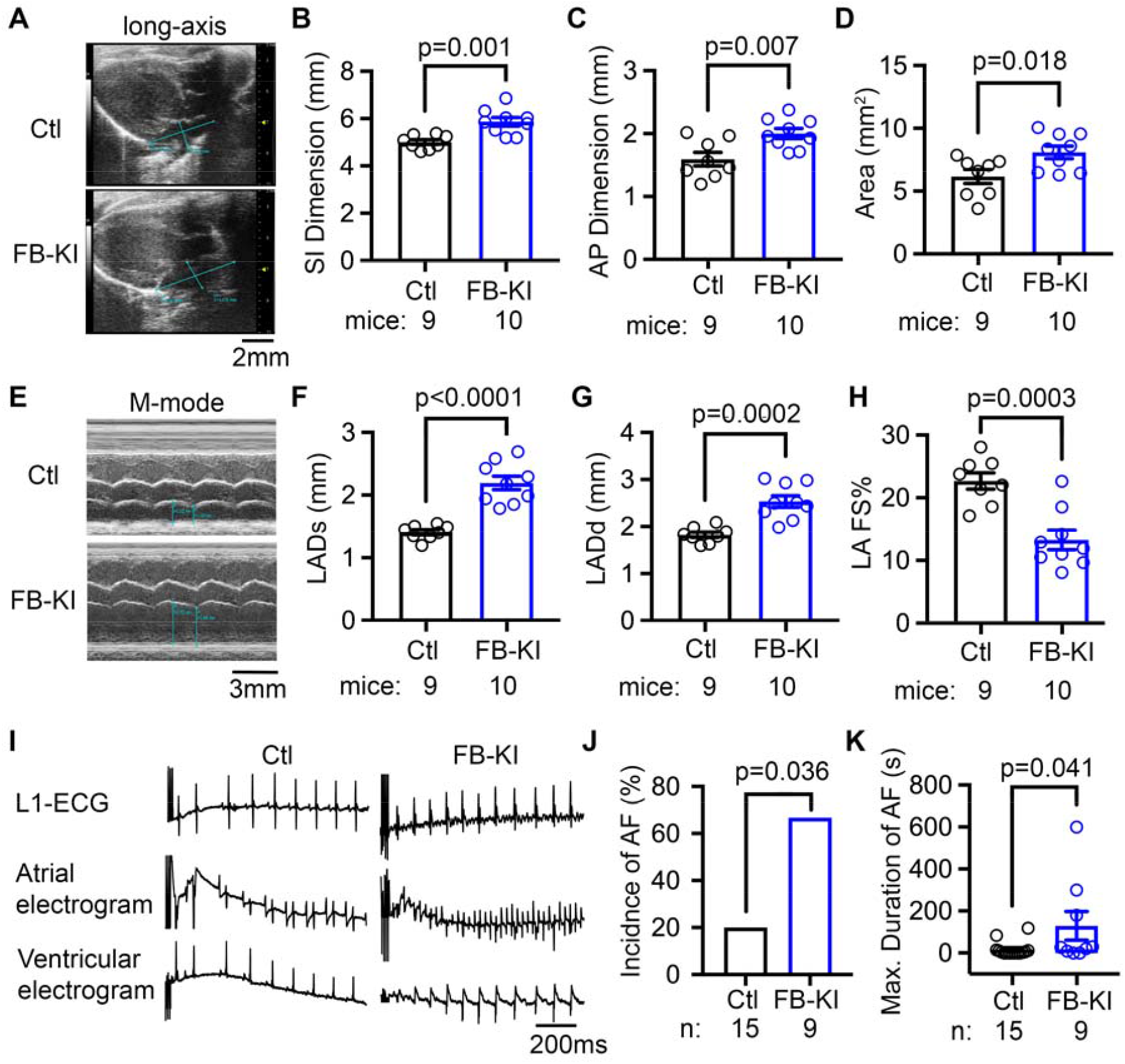
FB-specific activation of NLRP3 promotes atrial myopathy and enhances susceptibility to atrial arrhythmia. (**A**) Representative long-axis echocardiography images. (**B-D**) Quantifications of the superoinferior (SI, **B**) and anteroposterior (AP, **C**) dimensions and the area (**D**) of left atrium (LA). (**E**) Representative M-mode echocardiography of LA. (**F-H**) Quantifications of LA diameters (LADs, **F**; LADd, **G**) and fractional shortening (FS%, **H**). (**I**) Representative simultaneous recording of surface ECG (lead 1) and intracardiac electrograms showed sinus rhythm in Ctl and AF in FB-KI mice following rapid atrial pacing. (**J-K**) Incidence and the maximum duration of *in vivo* pacing-induced AF. p-values were determined with unpaired Student’s t-test (two-tailed) in **B, C, D, F, G**, and **H**, Fisher’s exact test in **J**, and Mann-Whitney test in **K**.

### FB NLRP3-inflammasome activation enhances fibroblast function and promotes fibrosis

To determine the consequences of FB-specific activation of NLRP3 inflammasome for cardiac FB function, we isolated primary FBs from Ctl and FB-KI mice. After 24-hour culture, the cardiac FBs of FB-KI mice exhibited an increased signal for αSMA staining (a marker of myofibroblast differentiation, **Figure 4A-B**). Consistently, the gel contraction assay revealed that the cardiac FBs of FB-KI mice exhibited stronger contractility than the cardiac FBs of Ctl mice with or without the TGF-β treatment (**Figure 4C-D**). Additionally, the scratch assay revealed that the cardiac FBs of FB-KI mice were more migratory (**Figure 4E**), and the percentage of Edu+ cells was also higher in FB-KI group than in Ctl group (**Figure 4F**), indicating increased proliferative activity. These results indicate that the function of cardiac FBs is enhanced in FB-KI mice. At the tissue level, Picrosirius red staining revealed that FB-KI mice exhibited a greater collagen deposition in both atria (p=0.014, **Figure 4G-H**) and ventricles (p=0.018, **Figure 4G, I**). Western blotting showed that pro- and mature collagen 1a (Col1a1) protein levels and profibrotic markers matrix metallopeptidase 9 (MMP9) and vimentin-1 were upregulated in FB-KI mice (**Figure 4J-K**). These results establish that FB-specific activation of NLRP3 inflammasome promotes the transdifferentiation of FBs to myofibroblasts, enhances migration and proliferation of FBs, and leads to cardiac fibrosis, a common pathogenic factor promoting atrial myopathy and AF ^2^.

**Figure 4.**
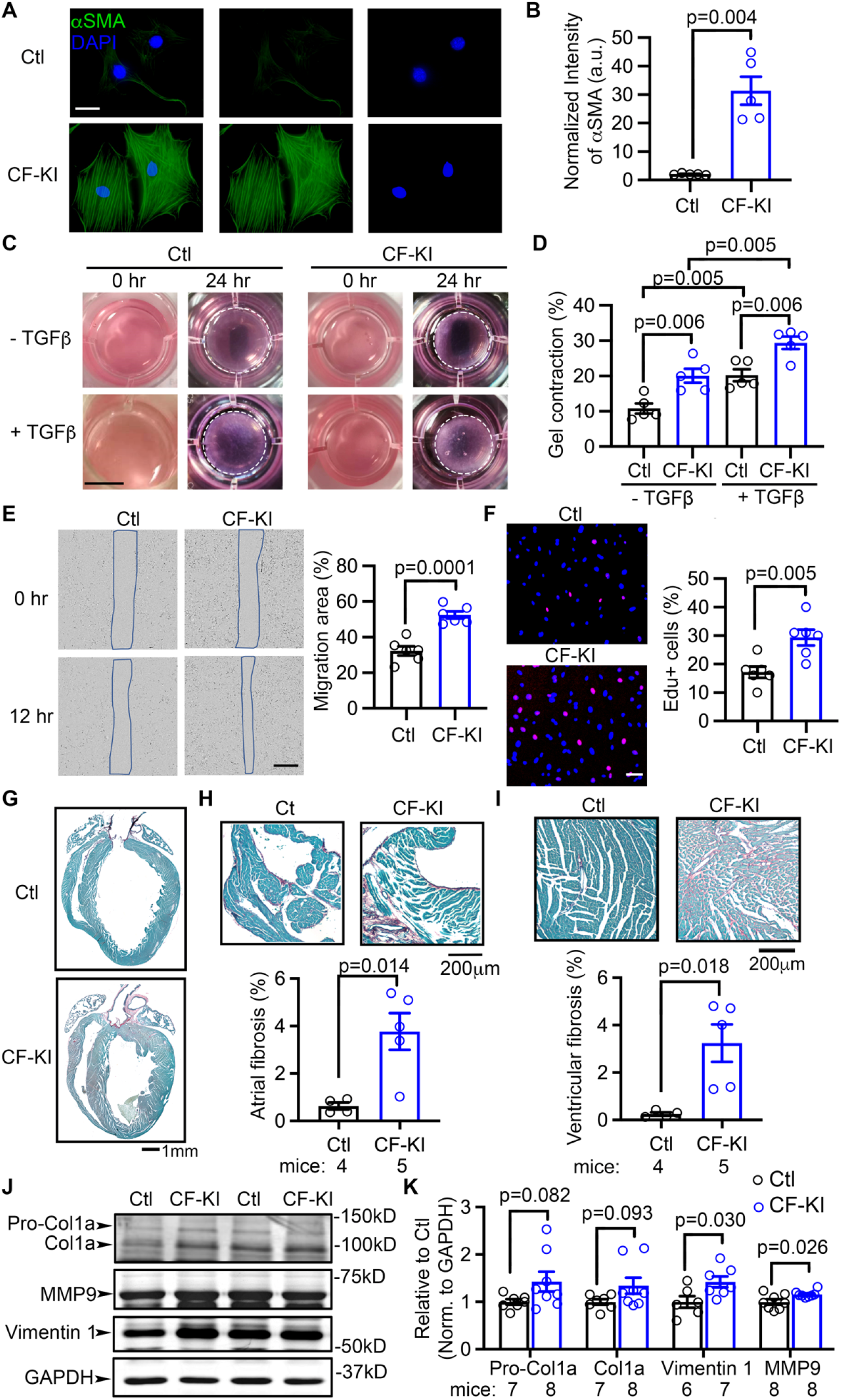
FB-specific activation of NLRP3 promotes cardiac fibrosis. (**A**) Representative images of αSMA staining and (**B**) quantification of αSMA intensity in isolated cardiac fibroblasts of Ctl and FB-KI mice, after 48 hours’ culture. (**C**) Representative images of gel contraction assay in isolated FBs with and without TGFβ challenge. White dash lines outlined the edges of gels after 24 hours’ culture. (**D**) Quantification of the contraction of gels (%), co-cultured with FBs of Ctl and FB-KI mice. (**E**) Representative images of scratch assay with the FBs of Ctl and FB-KI (left) and quantification of migration area (right). (**F**) Representative images of Edu staining (pink) in culture FBs of Ctl and FB-KI (left) and quantification of the Edu+ cells normalized to total cells. (**G**) Representative images of Picrosirius Red staining in whole heart. (**H**) Representative images of Picrosirius Red staining in atria and quantification of atrial fibrosis. (**I**) Representative images of Picrosirius Red staining in ventricles and quantification of ventricular fibrosis. Red color indicated the fibrosis region. (**J**) Representative Western blots and (**K**) quantifications of collagen 1a (Col1a) and fibrogenic proteins in atria of Ctl and FB-KI mice. p-values were determined with unpaired Student’s t-test (two-tailed) in **B, E, F, H, I** and **K**, and one-way ANOVA with Sidak’s multiple comparison in **D**.

### FB NLRP3-inflammasome activation alters cellular response

To determine how FB-specific activation of NLRP3 affects FB signaling, as well as the function and responses of other cell types in the heart leading to AF, we conducted snRNA-profiling using samples from Ctl and FB-KI mice 1-month post tamoxifen injection (**Figure 5A**). Following quality control and removal of low-quality nuclei, integrated cluster analysis on 22,840 nuclei across 2 samples (10,930 from Ctl, and 11,910 from FB-KI) was performed (**Figure 5B**). Based on known cell markers^20, 23^, we identified 16 clusters of cells including 3 clusters of fibroblasts (FB1, FB2, FB3), 1 cluster of myofibroblasts, and 3 clusters of CMs (CM1, CM2, CM3) (**Figure 5C**). Among the 3 clusters of FBs, the FB2-cluster expressed classical fibrotic markers, such as *Cilp* and *Postn*. The relative distribution of nuclei in each cluster was similar between Ctl and FB-KI mice, except for the CM1-, CM2-, and Epithelium-clusters (**Figure 5D**). We then performed GSEA pathway enrichment analyses based on the detected genes (FB-KI vs Ctl) in all clusters. The most enriched pathways across all clusters are related to metabolic activity, including ‘Oxidative Phosphorylation’, ‘Adipogenesis’, and ‘Fatty Acid Metabolism’ (**Figure 5E**). These data suggest that cardiometabolic stress, a contributor to atrial myopathy^27^, might be promoted by the constitutive activation of the NLRP3 inflammasome in FBs.

**Figure 5.**
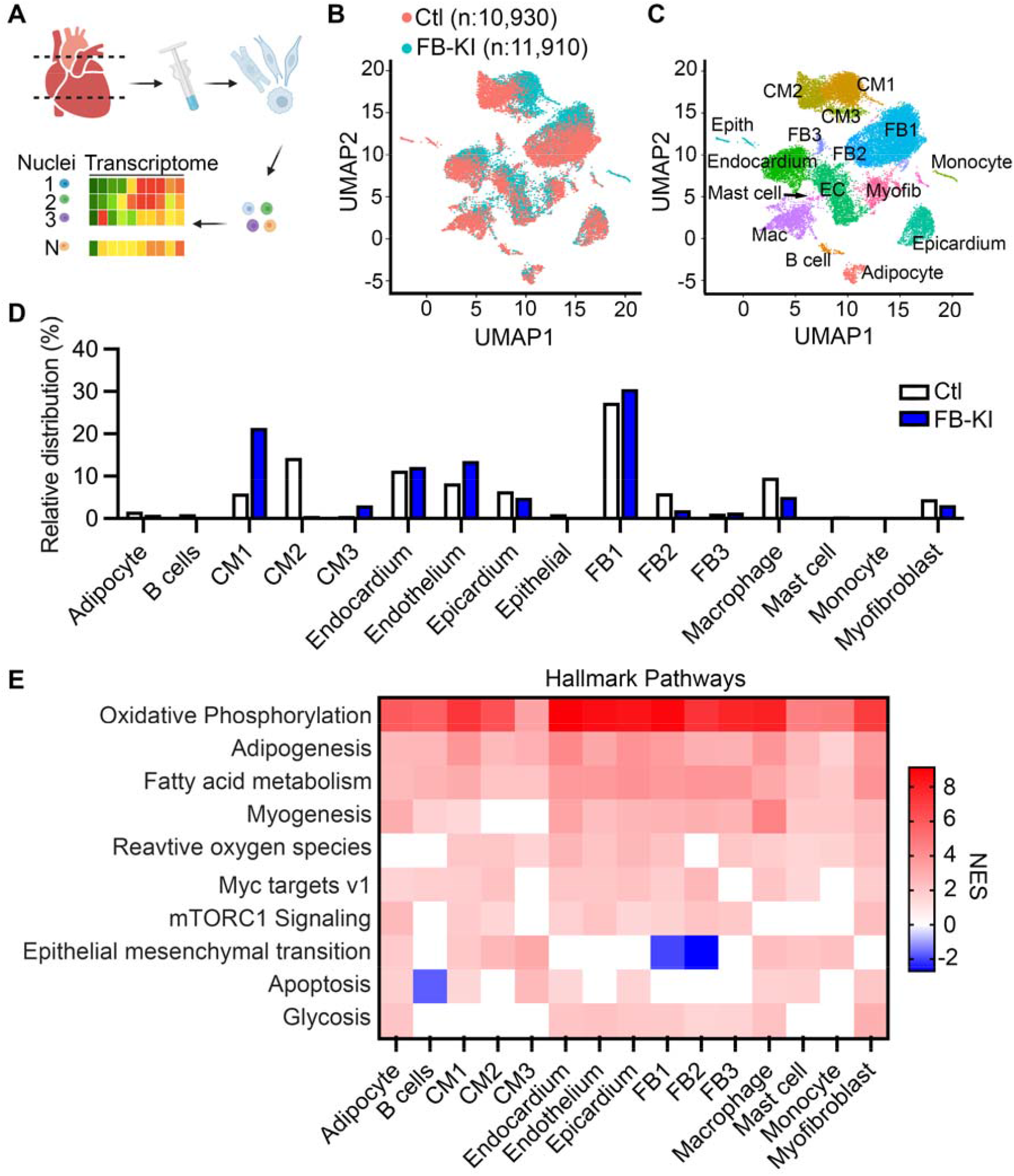
Single nuclei RNA profiling. (**A**) Experimental outline to profile the transcriptome of single nuclei. (**B**) UMAP representation of all filtered nuclei identified by snRNA-seq. (**C**) UMAP representation of 16 cell clusters identified by the known signature genes. CM, cardiomyocyte; FB, fibroblast; EC, endothelium; Epith, epithelium; Mac, macrophage; Myofib, myofibroblast. (**D**) Relative distribution of cell types in Ctl and FB samples. (**E**) Top 10 enriched Hallmark pathways using Gene Set Enrichment Analysis (GSEA) across 16 clusters of cells. NES, Normalized Enrichment Score.

Next, to assess how FB-specific inflammasome activation affects inter-cellular communications, we utilized the CellChat database^26^ and developed the ligand-receptor interaction networks for each mouse heart model (**Figure 6A**). Although FB2 was not the most abundant cluster of cells, the weighted interaction strength (reflected as the line thickness) initiated by the FB2-cluster was strongest among all clusters, suggesting that the FB2-initiated cell-cell communications were most prevalent in both Ctl and FB-KI samples. In addition, the comparison of the interaction maps between Ctl and FB-KI samples revealed that the weighted interactions were largely reduced (blue colored cells) in FB-KI with a few exceptions (red colored cells) (**Figure 6B**). Notably, Myofibroblast-cluster as a target received increased (FB-KI vs Ctl) in-coming interactions from all other clusters.

**Figure 6.**
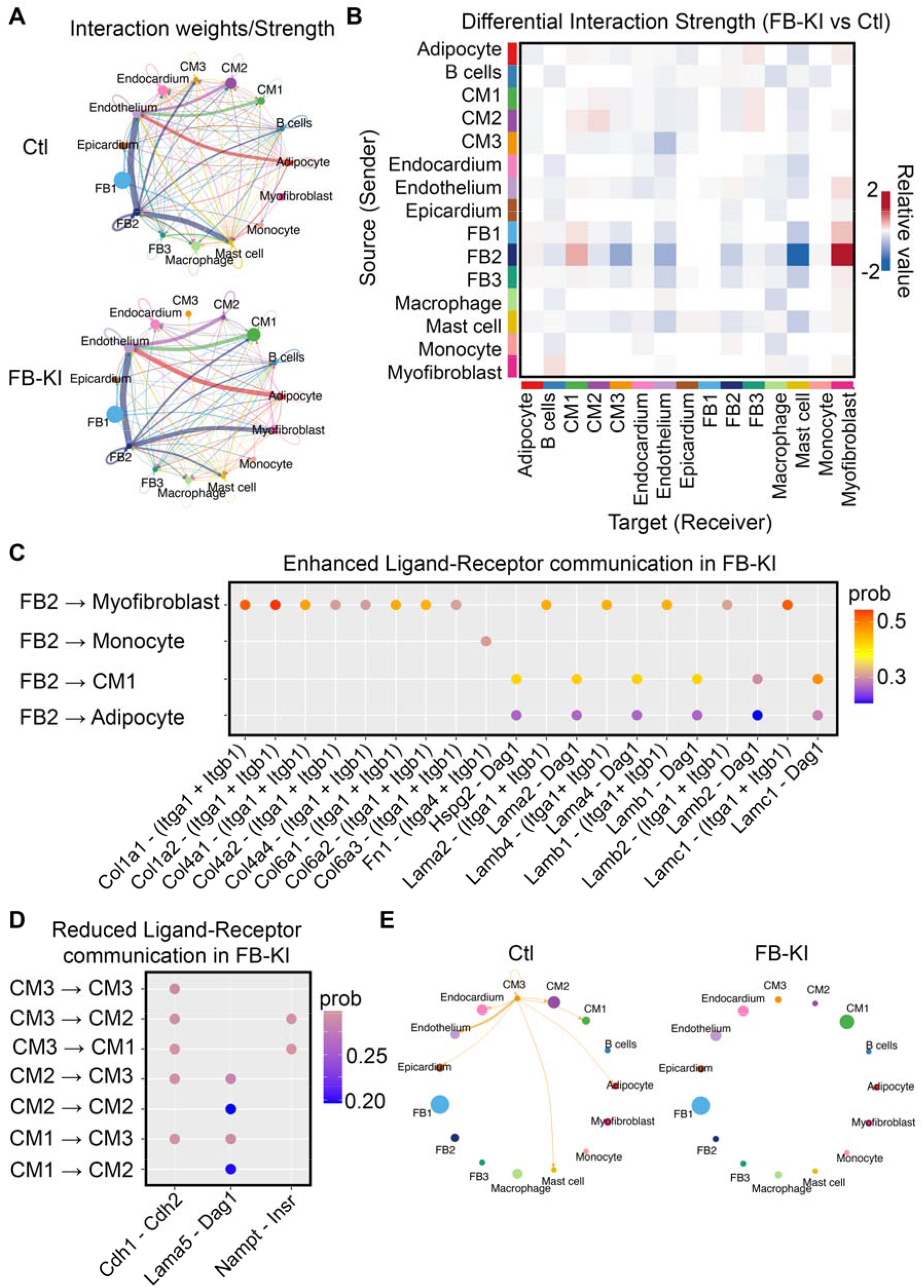
FB-specific activation of NLRP3 alters intercellular communication networks. (**A**) Weighted interaction networks in Ctl and FB-KI samples. (**B**) Differential interaction strength map between FB-KI and Ctl. Blue color indicates reduced communication. Red color indicates enhanced communication. (**C**) Enhanced ‘Ligand-Receptor’ communications in FB-KI mice initiated by the FB2-cluster. (**D**) Reduced ‘Ligand-Receptor’ communications in FB-KI mice among 3 CM clusters. (**E**) The complete loss of the CM3-initiated communications in FB-KI mice.

To gain insights into the altered interactions driven by the FB2 cluster (**Figure 6C**), we then extracted ligand-receptor interaction information. In terms of signaling pathways, the interactions between collagen 1/4/6 (*Col1/4/6*) and integrin α1/β1 (*Itga1/Itgb1*) were enhanced most between FB2 and Myofibroblast clusters, and the interactions between laminin α2/α4/β1/γ1 (*Lama2/a4/b1/c1*) and dystroglycan 1 (*Dag1*) or integrin α1/β1 (*Itga1/Itgb1*) were enhanced most between FB2 and CM1 clusters (**Figure 6C**). These results are consistent with the development of extracellular matrix remodeling in FB-KI hearts. Conversely, CM interactions were largely disrupted (**Figure 6B,D**). The reduced communications between CMs were mostly mediated through either reduced cell-cell adhesion between Cadherin 1 (*Cdh1*) and Cadherin 2 (*Cdh2*) or an impaired signaling transduction between laminin α5 (*Lama5)* and dystroglycan 1 (*Dag1*) or between nicotinamide phosphoribosyltransferase (*Nampt*) and insulin receptor protein (*Insr*) (**Figure 6D**). The CM3-initiated interactions were completely absent in FB-KI heart (**Figure 6E**). These results reveal that activation of FB NLRP3-inflammasome disrupts inter-cellular communication, probably because of increased FB activation with the development of endomysial fibrosis.

### FB NLRP3-inflammasome activation impairs atrial impulse conduction

Because the inter-cellular communications among CMs were impaired in FB-KI mice, we then determined whether and how this would affect electrophysiology at tissue level by optical mapping studies (**Figure 7A**). We found that conduction velocity (CV) was significantly reduced by about 40% (**Figure 7B**), while APD (at 20%, 50%, 70% and 90% of depolarization, **Figure 7C**) and AERP remained unchanged in FB-KI mice comparing with Ctl mice. We previously have established that CM-specific activation of the NLRP3 inflammasome in the CM-specific NLRP3 knockin mouse (CM-KI) causes a reentry-promoting abbreviation of APD and AERP^9^. These results indicate that selective activation of cardiac FB NLRP3 inflammasome in FB-KI mice does not affect refractoriness but impairs atrial conduction, creating a different reentrant substrate for AF in contrast to the CM-KI mice. The *ex vivo* heart rate was also reduced in FB-KI mice (**Figure 7D-E**). In addition to the enhanced fibrosis, known to reduce CV, we also found that the expression levels of the gap junction protein connexin 43 (Cx43) and fractional phosphorylation of Cx43-S368 proteins were reduced in atrial tissues of FB-KI mice (**Figure 7F**), while the protein levels of Nav1.5 (α-subunit of the cardiac Na^+^-channel) were unchanged and the Cx40 protein levels were increased in atria of FB-KI mice. Moreover, immunostaining revealed that the laterization of Cx43 was greater in atrial tissue (**Figure 7G**) and isolated atrial CMs (**Figure 7H**) of FB-KI mice. The ratio of Cx43 located at the lateral membrane over the Cx43 at the ID-region was significantly higher in atrial CMs of FB-KI mice than in Ctl mice (**Figure 7I**).

**Figure 7.**
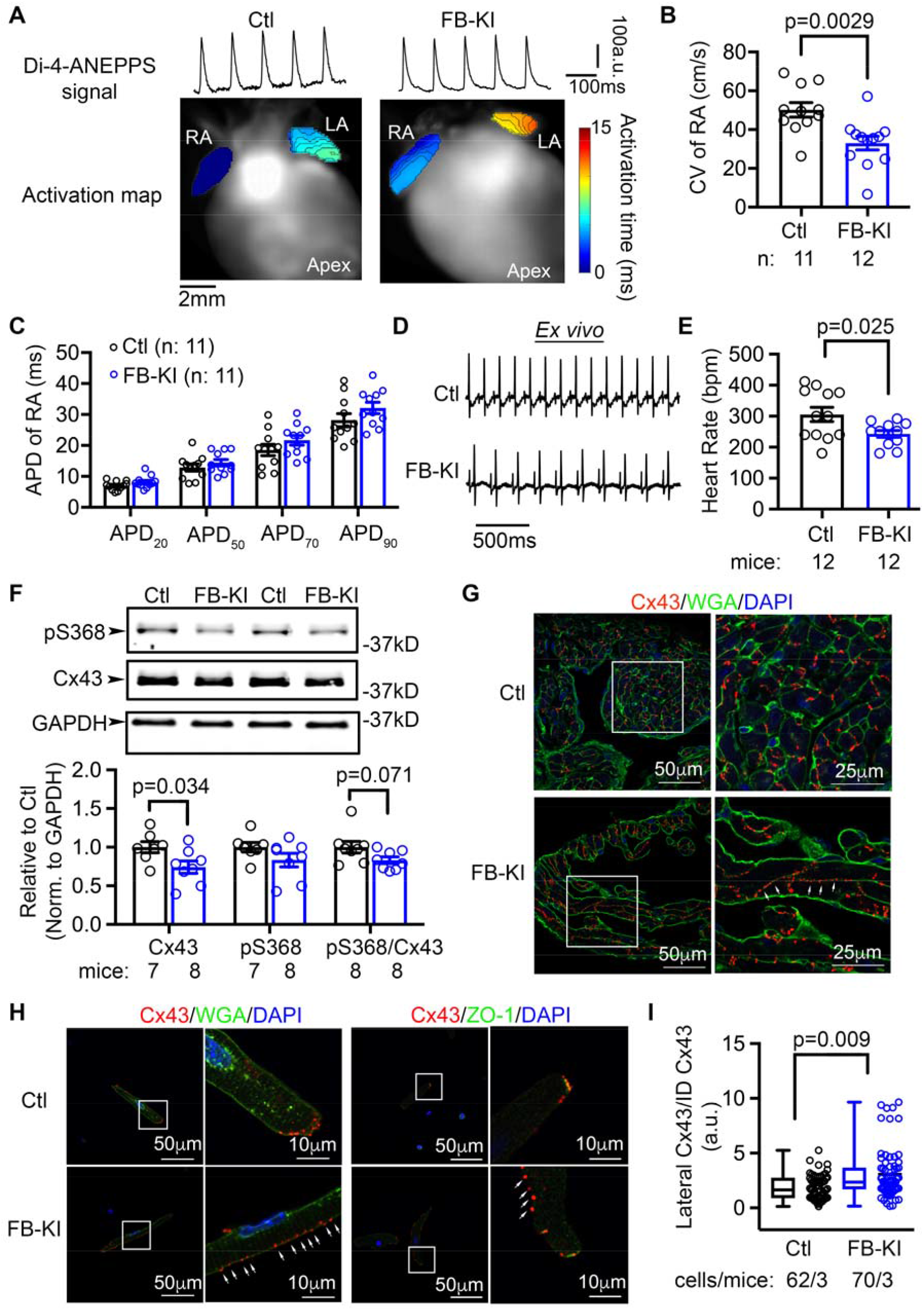
FB-specific activation of NLRP3 promotes a pro-arrhythmic substrate. (**A**) Representative optical di-4-ANEPPS signal and activation maps. (**B**) Conduction velocity (CV) in right atrium of Ctl and FB-KI mice. (**C**) Action potential duration (APD) at 20%, 50%, 70%, and 90% repolarization in right atrium (RA) (**D**) Representative *ex vivo* ECG recording revealed slower heart rate FB-KI mice. (**E**) Summary of *ex vivo* heart rate. (**F**) Representative Western blots and quantifications of Cx43 and phosphorylated Cx43-S368 in atria. (**G**) Representative images of immunohistochemistry of Cx43 in atria of Ctl and FB-KI mice. (**H**) Representative images of co-immunostaining of Cx43/WGA or Cx43/ZO-1 in atrial CMs of Ctl and FB-KI mice. (**I**) Quantification of the ratio of lateral Cx43 to intercalated disc (ID) Cx43 in atrial CMs of Ctl and FB-KI mice. p-values were determined with unpaired Student’s t-test (two-tailed) in **B, E**, and **F**, and Mann-Whitney test in **I**.

## DISCUSSION

Using multiple systems including human, dogs and mice, our results show that the NLRP3-inflammasome system is activated in FBs in the context of AF. A novel mouse model with the FB-restricted activation of the NLRP3 exhibits atrial myopathy and a higher susceptibility to AF. To our knowledge, this study is the first model addressing the arrhythmogenic consequences of FB inflammatory signaling. Our results demonstrate that, without ischemic injury, direct activation of FB inflammasome promotes fibrosis and defective conduction, which may underlie the development of AF-promoting atrial myopathy.

### Cell autonomous function of NLRP3 inflammasome signaling in FBs

Many studies have demonstrated that a variety of signaling systems in FBs, including Smad pathway, mitogen-activated protein kinases, reactive oxygen species, and Janus kinases, are important regulators of cardiac fibrosis^28^. Using the NLRP3 FB-KI model we are able to study the cell-autonomous effects of the cardiac FB NLRP3 inflammasome pathway activation. Our data demonstrate that the activation of FB NLRP3 inflammasome enhances the function of isolated cardiac FBs, evidenced as increased transdifferentiation, migration, and proliferation. snRNA-seq analysis reveal that the myofibroblasts of FB-KI mice are much more active in receiving incoming interactions initiated by other clusters of cells. The communication between the fibroblast FB2-cluster and the myofibroblast-cluster was enhanced most in FB-KI mice, which was primarily driven by the ‘laminin – integrin’ and ‘collagen – integrin’ pathways. These results validate the cell autonomous effect of the FB inflammasome pathway in modulating the function of cardiac FBs, and that the activation of NLRP3-inflammasome signaling in FBs is instrumental in the phenotypic switch from fibroblast to collagen-secreting myofibroblast^12^.

### FB abnormalities and gap junction remodeling

Based on snRNA-seq analysis, our data provide the first evidence that activation of FB NLRP3-inflammasome signaling can disrupt the communications and cell-cell contact among CMs. Consistently, we also show that CV was impaired in FB-KI, which is an established proarrhythmic substrate. The reduced CV in FB-KI model is attributed to gap junction remodeling, as a result of altered phosphorylation status and the cellular localization of Cx43. A reduction in Cx43 S368-phosphorylation is known to decrease the conductance of Cx43-based gap junctions ^29, 30^, while increased Cx43 lateralization reduces the expression of cell-coupling Cx proteins in gap junctions at the ID-region. Lateralized Cx43 may also form hemichannels with FBs and perhaps CMs, which might further enhance the heterogeneity of conduction and thus arrhythmogenicity ^31^. We also noted increased expression of Cx40, along with reduced expression of Cx43; an increase in the Cx40/Cx43 ratio has previously been shown to slow atrial propagation in synthetic strands of atrial CMs ^32^. How the FB-specific activation of NLRP3-inflammasome affects the protein level of Cx43 and Cx40 and the phosphorylation of Cx43 requires future investigations.

### Inflammatory signaling in FBs and CMs promotes AF via distinct but complementary arrhythmogenic mechanisms

Although CMs present the major cell population in the heart, the role of FBs in cardiac electrophysiology and AF pathogenesis has attracted increasing interests in recent years, partially because fibrosis is an important contributor to AF pathophysiology ^28, 33, 34^. We previously have shown that the activation of NLRP3 in CMs enhances AF susceptibility by promoting abnormal Ca^2+^-release events in CMs along with AERP-shortening (due to increased expression of the ultra-rapid rectifier K^+^-channel) in NLRP3 CM-KI mice ^9^. In contrast to the major phenotypes observed in CM-KI mice, the current study reveals that the activation of NLRP3 inflammasomes in resident FBs enhances atrial arrhythmogenesis by promoting connexin dysfunction and extracellular matrix remodeling in NLRP3 FB-KI mice, two crucial contributors to the development of the conduction abnormalities. Thus, our studies suggest that the activation of the cardiac NLRP3-inflammasome systems in CMs and cardiac FBs creates unique (triggered activity and abbreviated refractoriness in CMs and connexin and fibrotic remodeling in cardiac FBs) alterations, which likely combine to promote clinical AF in patients. Therefore, targeting NLRP3 in a non-cell type specific manner may be a more effective anti-AF treatment strategy, because it is expected to modulate multiple proarrhythmic mechanisms.

### Study limitations

Because of the very small number of FBs that can be isolated from atrial biopsies from patients, we were unable to assess the protein levels of the NLRP3-inflammasome system in atrial FBs of AF patients. We were, however, able to assess NLRP3 system protein changes in a canine model of AF-associated remodeling and identified clear changes. Due to technical challenges, we were unable to yield enough atrial fibroblasts from mice for Western blot and fibroblast functional assays. Therefore, the murine FB-samples are mixture of atrial FBs and ventricular FBs. Whether atrial FBs or ventricular FBs are more prone to inflammasome activation and fibrosis remodeling without ischemic injury requires further investigation. In our FB-KI mouse model, the NLRP3-inflammasome system was activated in the Tcf21-expressing resident FBs. The role of NLRP3 inflammasome in cardiac FBs derived from other precursors such as myeloid and the endogenous mesenchymal stem cells remain to be determined ^35, 36^. The discovery of enhanced metabolic pathways across different cell populations in FB-KI mice is interesting. However, it is beyond the scope of the current study to determine whether and how FB inflammasome activation modulates mitochondrial metabolism in the different cardiac cell types.

## Conclusions

Our data reveal that the FB-restricted activation of NLRP3 inflammasome promotes the concomitant development of atrial myopathy and AF. Activation of NLRP3 inflammasome in resident FBs exhibits cell autonomous function by enhancing the transdifferentiation, migration, and proliferation of cardiac FBs, subsequently causing fibrosis, connexin remodeling, and an arrhythmogenic substrate for AF that differs from the atrial arrhythmogenic consequences of CM-restricted activation of NLRP3. This study establishes the NLRP3 inflammasome as a novel FB-signaling pathway contributing to arrhythmogenic myopathy.

